# CNBP controls c-Rel dependent IL-12β gene transcription and Th1 immunity

**DOI:** 10.1101/258731

**Authors:** Yongzhi Chen, Shruti Sharma, Patricia A. Assis, Zhaozhao Jiang, Andrew J. Olive, Saiyu Hang, Jennifer Bernier, Jun R. Huh, Christopher M. Sassetti, David M. Knipe, Riccardo T. Gazzinelli, Katherine A. Fitzgerald

## Abstract

An inducible program of inflammatory gene expression is a hallmark of antimicrobial defenses. Herein, we identified Cellular nucleic acid binding protein (Cnbp) as a specific regulator of interleukin-12β gene transcription and Th1 immunity. Cnbp resides in the cytosol of macrophages and translocates to the nucleus in response to a broad range of microbial ligands. Cnbp-deficient macrophages had a selective impairment in their ability to induce IL12β gene transcription. Cnbp interacted with c-Rel, an NFκB/Rel family member that controls IL12β gene transcription. c-Rel nuclear translocation and DNA binding activity were dependent on Cnbp. Furthermore, Cnbp itself bound the IL12β promoter. Lastly, Cnbp-deficient mice were more susceptible to acute toxoplasmosis associated with reduced production of IL12β, as well as a reduced Th1 cell IFNγ response essential to control parasite replication. Collectively, these findings identify Cnbp as a key regulator of c-Rel dependent IL12β gene transcription and Th1 immunity.

## Introduction

The acute inflammatory response is induced as a first line of defense against pathogen invasion. Innate immune cells including macrophages and dendritic cells (DCs) are the primary drivers of acute inflammation. These processes are initiated when myeloid cells recognize pathogen-associated molecular patterns (PAMPs) via pattern recognition receptors (PRRs). A variety of PRRs have been identified including Toll-like receptors (TLRs), Nod-like receptors, Aim2-like receptors, Rig-I-like receptors, and C-type lectins (Cao, 2016; Chen et al., 2016; O’Neill et al., 2013; Thompson et al., 2011; Trinchieri and Sher, 2007). This PRR–PAMP interactions in turn promote the activation of signal transduction pathways that converge on transcription factors resulting in changes in expression of hundreds of genes involved in antimicrobial defense, phagocytosis, cell migration, metabolic reprogramming, tissue repair, and regulation of adaptive immunity(Liu et al., 2016; Medzhitov and Horng, 2009; Trinchieri and Sher, 2007). Signal-dependent activation of transcription factors such as NFκB, Interferon Regulatory factors (IRFs) and Signal transducers and activators of transcription (STAT) together with transcriptional co-regulators, and chromatin-modifying factors collaborate to control these transcriptional programs (Ghosh and Hayden, 2008; Smale and Natoli, 2014). The timing and duration of these responses must be tightly controlled during infection to ensure appropriate and measured immune cell activation and anti-microbial defenses. Inappropriate regulation of these processes leads to chronic inflammation and diseases such as autoimmunity, atherosclerosis and cancer.

The proinflammatory cytokine interleukin-12 (IL-12) acts as a bridge between the innate and the adaptive immune response. Originally called NK cell stimulatory factor or CTL maturation factor, IL-12 is unique among the proinflammatory heterodimeric cytokines consisting of the two subunits p40 and p35 (Murphy, 2009; Teng et al., 2015; Trinchieri, 1995). Products from microorganisms including bacteria, intracellular parasites, fungi, double-stranded RNA, bacterial DNA and CpG-containing oligonucleotides are potent inducers of IL-12 in macrophages, monocytes, neutrophils and dendritic cells(Mason et al., 2002). The induction of IL-12 is critical for host resistance against different categories of pathogens including Cytomegalovirus (Orange et al., 1995), *Mycobacterium tuberculosis* (Mtb) (Cooper et al., 2002; Holscher et al., 2001; Khader et al., 2006), *Listeria (Tripp et al., 1993), Leishmania (Sypek et al., 1993), Toxoplasma (Dupont et al., 2015; Gazzinelli et al., 1993; Gazzinelli et al., 2014; Sher et al., 2003)* and *Salmonella* (Lehmann et al., 2001; Schulz et al., 2008) through the establishment of T helper 1 cell and IFNγ mediated immune responses (Biron and Gazzinelli, 1995; Gazzinelli et al., 2014).

The inducible expression of IL12 is regulated at the level of gene transcription (Sanjabi et al., 2000; Smale, 2012). The mouse IL12β promoter contains several cis-elements that bind a number of transcription factors including NFκB/Rel family members, IRF5 and CCAAT/enhancer-binding protein (C/EBP)(Bradley et al., 2003; Koshiba et al., 2013; Sanjabi et al., 2000; Sanjabi et al., 2005). The NFκB/Rel family of transcription factors includes the proteins RelA (p65), RelB, c-Rel, NFkBI (p105/p50), and NFκB2 (p100/p52)(Caamano and Hunter, 2002; Dev et al., 2011; Hayden and Ghosh, 2011). These proteins can homo- and heterodimerize possessing unique specificities in regulating target gene expression. Rel binding sites have been identified in the promoters of cytokine and immune effectors in macrophages. Many proinflammatory genes induced downstream of TLRs and other PRRs engage the RelA/p50 heterodimeric complex to control their expression. In the case of IL12, however, c-Rel has been defined as the critical NFκB family member responsible (Mason et al., 2002; Sanjabi et al., 2000).

Cellular nucleic acid binding protein (Cnbp), also called zinc-finger protein 9 (ZNF9), is a highly conserved zinc-finger protein with seven tandem repeats of CCHC zinc finger knuckles and one arginine-glycine-glycine (RGG) box (Calcaterra et al., 2010). This DNA- and RNA-binding protein with broad sequence specificity has been associated with diverse cellular functions, including transcription and translation (Benhalevy et al., 2017; Chen et al., 2013; Margarit et al., 2014). Cnbp is linked to age-related sporadic inclusion body myositis (sIBM), an inflammatory muscle disease characterized by progressive muscle weakness and atrophy (Niedowicz et al., 2010). Furthermore, a dominantly transmitted (CCTG)n expansion in intron 1 of the Cnbp gene leads to myotonic dystrophy type 2 (DM2)(Meola and Cardani, 2015; Raheem et al., 2010; Sun et al., 2011; Thornton et al., 2017), a disease associated with a high frequency of autoantibodies and autoreactive T cells (Tieleman et al., 2009). These observations suggest that Cnbp may have some function in the immune system.

Here we identify Cnbp as a cytosolic protein that moves to the nucleus in an inducible manner where it acts as a highly specific regulator of the inflammatory program. We generated mice lacking Cnbp and found that macrophages lacking Cnbp were compromised in their ability to induce IL-12 p40 (IL12β) gene expression, at early phases during the response to DNA sensing pathways as well as TLR ligands and viruses which signal via RIG-I. Importantly, Cnbp was not a broad regulator of the NFκB/Rel family of transcription factors. Rather, Cnbp specifically controlled the activity of c-Rel. c-Rel but not RelA (p65) nuclear translocation was dependent on Cnbp. Furthermore, we provide *in vivo* evidence for the importance of the Cnbp-c-Rel-IL12 pathway in controlling acute Toxoplasmosis *in vivo*. Collectively, these studies reveal a previously unrecognized role for Cnbp as a novel transcriptional regulator controlling the c-Rel-IL-12-Th1 axis.

## Results

### Identification of CNBP a cytosolic dsDNA binding protein

To systematically identify proteins involved in the recognition of foreign nucleic acids, we took advantage of immune stimulatory oligonucleotides (ATr-ODN) that we had previously shown were potent stimulators of inflammatory cytokines via the STING pathway (Gallego-Marin et al., 2018; Sharma et al., 2011). In an effort to identify receptors or regulators of dsDNA induced signaling pathways, ATr-ODN and a non-stimulatory ODN (ATr-AA ODN) were used in pull down experiments to capture DNA binding proteins from macrophage cytosolic extracts. ATr-ODN binding proteins were eluted and separated by SDS PAGE. Immunoblotting using antibodies to known components of the cytosolic DNA sensing pathway identified STING, DDX3x, as well as TBK1 in these pull downs (Supplemental Fig. 1A). To identify additional components of these pathways we also performed liquid chromatography mass spectrometry (LC-MS) on these ATr-ODN pulldowns. The most enriched of the proteins identified in the ATr-ODN but not the non-stimulatory ATr-AA-pulldowns was Cellular nucleic acid binding protein (Cnbp) (Supplementary Fig 1B). Cnbp also called Zinc finger protein 9 is a highly conserved zinc-finger protein with seven tandem repeats of CCHC zinc finger knuckles and one arginine-glycine-glycine (RGG) box. Cnbp mRNA is constitutively expressed in numerous tissues but particularly enriched in the spleen, lung and muscle (Fig. 1A). Immunofluorescence staining of endogenous Cnbp protein in macrophages showed that Cnbp was predominantly localized in the cytoplasm at steady state (Fig. 1B). This is consistent with our ability to isolate this protein from cytosolic extracts of macrophages.

**Figure 1.**
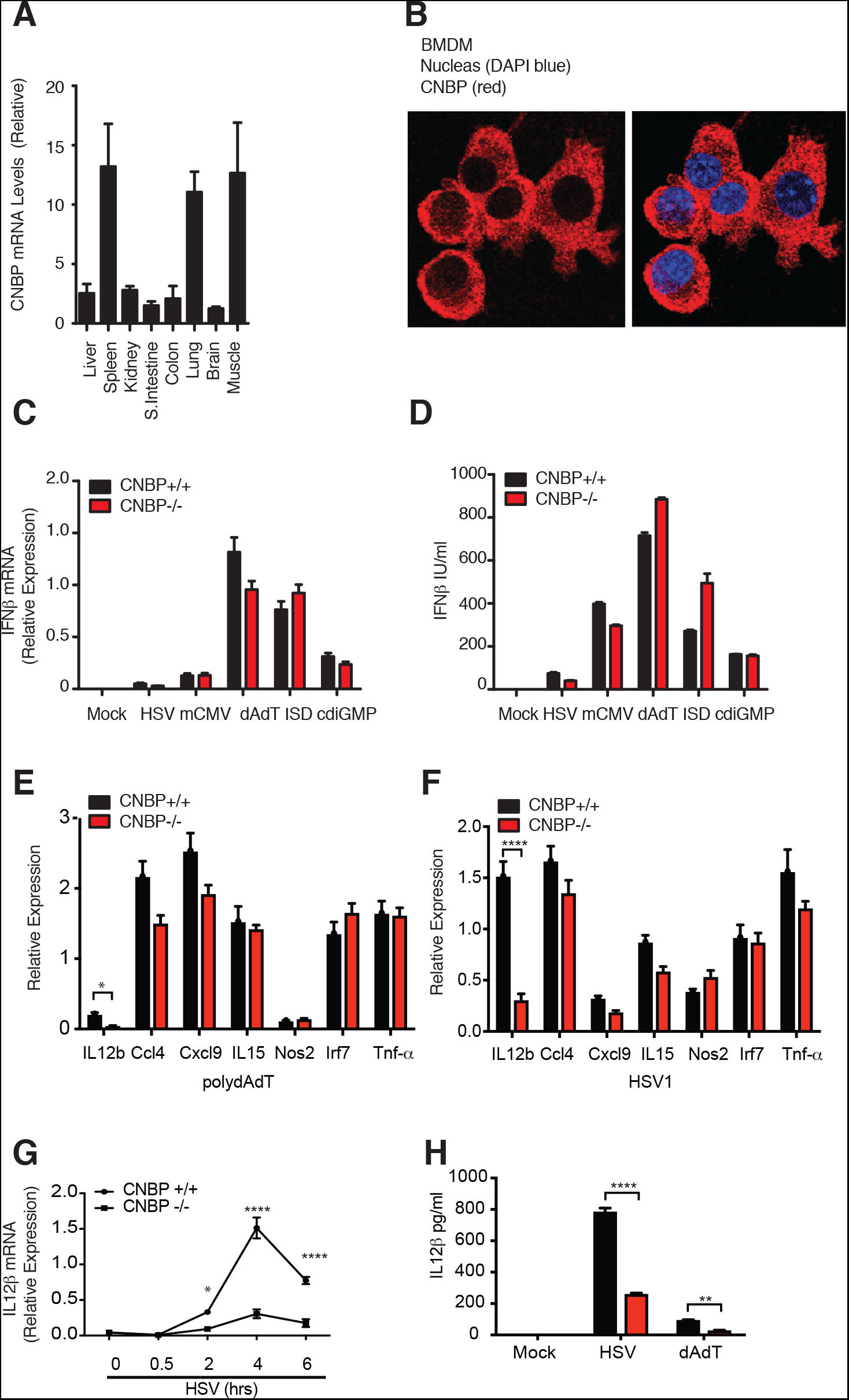
Identification of CNBP a cytosolic dsDNA binding protein and regulating the cytosolic dsDNA induced IL12p40 response. (A) Tissue-based expression of Cnbp in mouse (normalized to GAPDH) N=3. (B) Confocal microscopy of primary BMDMs probed with anti-CNBP. (C and D) IFNβ mRNA and protein level in CNBP+/+ and CNBP−/− BMDMs treated with DNA ligands including HSV, mCMV, dAdT, ISD or cdiGMP were detected through qRT-PCR (C) and ELISA (D) (E and F) qRT-PCR analysis of IL12β, Ccl4, Cxcl9, IL15, Nos2, Irf7 or Tnf-α mRNA in Cnbp WT or KO BMDMs left unstimulated or transfected with PolydAdT (E) or HSV1 infection (F). (G) qRT-PCR analysis of IL12β in Cnbp WT or KO BMDMs mock infected or infected with HSV1 at different time points. (H) IL12β protein levels in CNBP+/+ and CNBP−/− BMDMs treated with HSV or dAdT were detected through ELISA. Error bars represent s.d. of triplicate technical replicates (C, E-G) or s.e.m. of triplicate biological replicates (A, D, H). Data are representative of one (A), two (B) or three (C–H) independent experiments. *p < 0.05, ****p < 0.0001. See also Figure S1.

### CNBP regulates the cytosolic dsDNA induced IL12p40 response

To investigate the contribution of CNBP to the cytosolic DNA sensing pathway in macrophages, we generated Cnbp-deficient mice. While Cnbp-deficient mice are viable, homozygous knockout animals were born at a lower than expected Mendelian frequency. Heterozygous mice were intercrossed resulting in 29.5% wild type, 55.5% heterozygous and 15% homozygous mice. Homozygous CNBP-knockout animals were runted at birth, they reached weights equivalent to their wild type littermate controls by 6 weeks. BMDM were generated from *Cnbp^+/+^* and *Cnbp^−/−^* littermate controls. Macrophage differentiation was examined by staining cells for CD11b and F4/80. Both *Cnbp^+/+^* and *Cnbp^−/−^*macrophages had equivalent F4/80 and CD11b staining patterns indicating normal macrophage differentiation (Supplemental Fig. 1C). There were also normal numbers of macrophages and dendritic cells in the peritoneum (Supplemental Figs 1D and E). Quantitative PCR (qPCR) and immunoblotting confirmed loss of Cnbp expression in bone marrow–derived macrophages (BMDMs), bone marrow–derived dentritic cells (BMDCs) and peritoneal macrophages (PECs) from Cbnp−/- mice as indicated (Supplemental Figs 1F and G). Lastly, numbers of CD4 and CD8 positive T cells were equivalent in the spleens of *Cnbp^+/+^* and *Cnbp^−/−^* littermate controls (Supplemental Figs 1H and I).

Type I IFN induction is a hallmark of the cytosolic DNA sensing pathways and we first investigated if Cnbp was involved in its production in macrophages. *Cnbp^+/+^* and *Cnbp^−/−^* macrophages were infected with Herpesviruses Herpes simplex virus (HSV-1-ICP0-deficient mutant) and mouse cytomegalovirus (mCMV) or stimulated with polydAdT, the immune stimulatory dsDNA ISD and cdiGMP all of which signal via the STING pathway and IFNβ levels measured. In all cases, IFNβ mRNA and protein levels were induced at similar levels between the genotypes (Fig. 1C and D). Consistent with this result, replication of mCMV and MHV-68 was unaffected in *Cnbp^−/−^* mouse embryonic fibroblasts (MEFs) compared with wild-type cells (Supplementary Figs 1J and K). We also measured TNFα mRNA and protein levels which were also comparable between *Cnbp^+/+^* and *Cnbp^−/−^* macrophages (Supplemental Fig. 1L) These results indicate that Cnbp is not a receptor for cytosolic dsDNA controlling the type I IFN or NFκB-dependent TNFα response.

We expanded this analysis to evaluate the inducible expression of a broader panel of immune response genes in cells exposed to polydAdT or HSV1. While the inducible expression of several genes including Ccl4, Cxcl9, IL15, Nos2, Irf7 and TNFα were induced normally in *Cnbp^+/+^* and *Cnbp^−/−^* macrophages, the only impact on the DNA sensing pathway we observed was on the induction of the IL12β gene (Fig 1E, F). *Cnbp^−/−^* BMDMs were impaired in their ability to induce IL12β gene transcription following infection with HSV1 or transfection with polydAdT. We confirmed these results by monitoring the kinetics of IL12β mRNA expression during infection with HSV1 from 0-6hrs (Fig 1G) and at the protein level (Fig 1H). Collectively, these results indicate that Cnbp was not involved in the DNA sensing STING-IRF3-IFNβ pathway but identified a role for Cnbp in controlling IL12β gene expression.

### CNBP is a broad regulator of the IL12β response in the TLR and RLR pathway

To broaden our investigation of Cnbp in the control of inflammatory responses, we next evaluated the role of Cnbp in the TLR signaling pathway. We used multiplex gene expression analysis (Nanostring) to compare the inducible expression of a panel of 254 inflammatory genes in Cnbp^+/+^ and Cnbp^−/−^ macrophages stimulated with LPS. These genes represent a broad range of inflammatory cytokines, type I interferons, interferon stimulated genes and other pathways that included apoptosis, EGF, interleukin signaling, Ras, T-cell receptor and Toll-like receptor signaling genes. While most of these LPS induced genes were induced at equivalent levels in Cnbp^+/+^ and Cnbp^-/-^ macrophages, the expression of IL12β was markedly reduced in Cnbp^-/-^ macrophages compared to control cells (Fig 2A). A heatmap depicting ~80 of the most highly inducible LPS regulated genes in wild type and KO cells is shown. In addition to this rather selective impairment of IL12β gene expression the levels of interleukin-10 mRNA were higher in *Cnbp^−/−^* cells relative to their wild type *Cnbp^+/+^* counterparts (Fig. 2A). We confirmed these effects for IL12β and IL10 using qRT-PCR (Fig. 2B, C). This effect was not restricted to BMDM as similar results were seen in peritoneal macrophages (PEC) and dendritic cells (BMDC) from Cnbp^−/−^ mice (Fig 2D, E). Similar to what we had observed for HSV1 and polydAdT the defect in IL12β gene expression was clear over a time course of LPS (Fig 2F). Moreover, the requirement for Cnbp in controlling IL12β gene expression was also observed in cells infected with Sendai virus, a paramyxovirus which signals via RIG-I (Fig 2G) and in cells stimulated with poly I:C, a TLR3 and MDA5 ligand (Fig 2H). Levels of IL12β and IL-10 protein measured by ELISA but not TNFα were reduced for all of these stimuli (Supplemental Fig 2A, B, C). Taken together, all of these data indicate that Cnbp regulates IL12p40 mRNA and production of IL12β in response to a broad range of PRR ligands.

**Figure 2.**
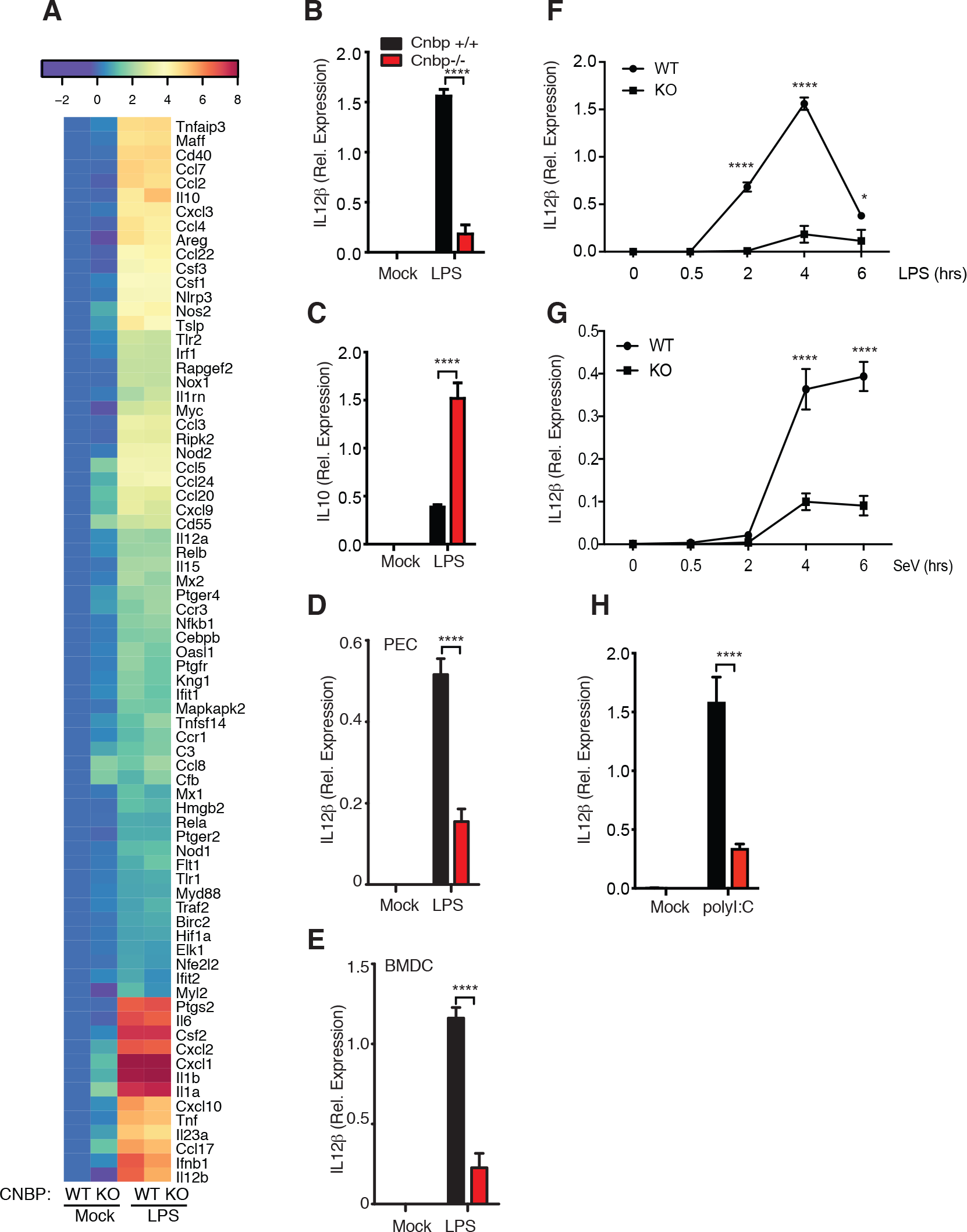
CNBP is a broad regulator of the IL12p40 response in the TLR and RLR pathway. (A) Heatmap of gene expression in WT and KO BMDMs treated or not treated with LPS and analyzed by NanoString. N=3 (B and C) qRT-PCR analysis of IL12β (B) and IL10 (C) in Cnbp WT or KO BMDMs left unstimulated or stimulated with LPS. (D and E) qRT-PCR analysis of IL12β mRNA in Cnbp WT or KO peritoneal macrophages (PEC) (D) or BMDCs (E) left unstimulated or stimulated with LPS. (F and G) qRT-PCR analysis of IL12β in Cnbp WT or KO BMDMs left unstimulated or stimulated with LPS (F) or SeV (G) at different time points. (H) qRT-PCR analysis of IL12β in Cnbp WT or KO BMDMs left unstimulated or stimulated with PolyI:C. Error bars represent s.d. of triplicate technical replicates (B-H). Data are representative of three independent experiments with similar results. *p < 0.05, ****p < 0.0001. See also Figure S2.

### CNBP translocates to the nucleus in response to PRR ligation

We next wanted to understand how Cnbp regulates IL12β gene expression and the molecular basis for the specificity of this response. As described in Fig 1, Cnbp is relatively abundant in myeloid cells, however we also found that Cnbp expression could be further upregulated in cells activated via TLR stimulation or virus infection. BMDMs were stimulated with LPS or infected with Sendai virus and the levels of Cnbp mRNA measured by qRT-PCR. This analysis revealed that Cnbp was upregulated after LPS stimulation or SeV infection (Fig. 3A). To further explore the mechanisms by which Cnbp was induced under these conditions, BMDMs were pretreated with DMSO (control) or BAY11-7082, an irreversible inhibitor of the IKK kinases, followed by LPS stimulation or SeV infection. The upregulation of Cnbp was abolished in BAY11-7082-treated BMDMs relative to the control cells (Fig. 3B), indicating that the inducible expression of Cnbp was dependent on the NFkB signaling pathway.

**Figure 3.**
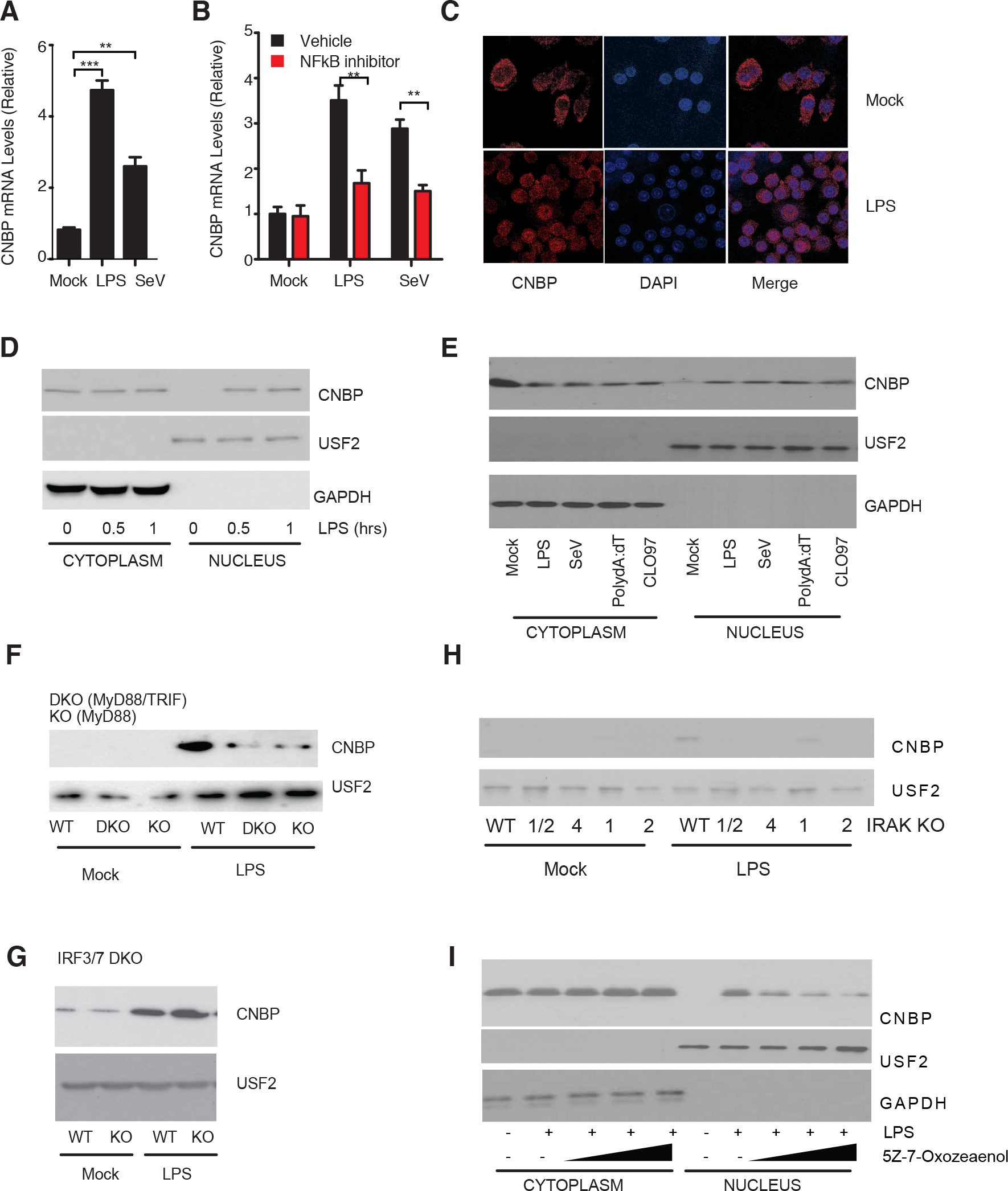
CNBP translocates to the nucleus in response to PRR ligation. (A) qRT-PCR analysis of Cnbp expression in BMDMs stimulated with LPS or SeV. (B) WT BMDMs were pretreated with the NF-kB inhibitor BAY11-7082 (10 µM) or DMSO, followed by LPS stimulation or SeV infection. (C) Confocal analysis of Cnbp distribution in primary macrophage stimulated with LPS; (D) The cytosolic and nuclear extracts were analyzed for Cnbp by Western blotting in WT BMDMs treated with LPS. Anti-GAPDH or anti-USF2 were used for the cytosolic or nuclear loading control respectively. (E) The cytosolic and nuclear extracts were analyzed for Cnbp by Western blotting in WT BMDMs treated with LPS, SeV, Poly dAdT or CLO97. (F-H) Nuclear extracts were analyzed for Cnbp by Western blotting in MyD88−/- and MyD88−/-TRIF−/- (F), IRF3/7−/- (G) or IRAK1/2−/-, IRAK4−/-, IRAK1−/-, IRAK2−/- (H) BMDMs treated with LPS. (I) The cytosolic and nuclear extracts were analyzed for Cnbp by Western blotting in WT BMDMs treated with the TAK1 kinase inhibitor 5Z-7 Oxozeaenol in the presence of LPS. Error bars represent s.d. of triplicate technical replicates (A-B). Data are representative of three independent experiments with similar results. **p < 0.01, ***p < 0.001. See also Figure S3.

Although Cnbp is inducible in macrophages it is expressed at steady state predominantly localized in the cytosol. We speculated that Cnbp may relocalize when macrophages are activated through TLRs or other PRRs. To test this, we used confocal microscopy to monitor Cnbp localization both at baseline and following exposure to LPS. Confocal microscopy revealed that Cnbp translocated from the cytosol to the nucleus in response to LPS (Fig 3C). We confirmed this by purifying nuclear extracts and monitoring Cnbp nuclear accumulation by immunoblotting. In resting cells, CNBP was exclusively cytosolic excluded from the nuclear compartment. Upon LPS stimulation, Cnbp translocated to the nucleus within 30 mins (Fig 3D). Cnbp also translocated to the nucleus in cells exposed to SeV, polydAdT or CL097, a TLR7 ligand indicating that multiple TLRs and other PRRs trigger Cnbp nuclear translocation (Fig 3E). These experiments were controlled by detecting USF2 and GAPDH, as proteins that are found exclusively in the nuclei and cytosol, respectively.

We next wanted to understand the pathway responsible for CNBP nuclear translocation. Following TLR4 activation, downstream signaling proceeds through the recruitment of TIR domain containing adapter molecules. To evaluate if nuclear translocation of Cnbp proceeded via TIR domain signaling we monitored Cnbp nuclear translocation in macrophages from mice lacking MyD88 and TRIF. These double KO mice fail to induce downstream TLR signaling (Yamamoto et al., 2003). The nuclear translocation of Cnbp was significantly reduced in macrophages from these DKO (Fig 3F). This effect was largely MyD88 dependent since macrophages from MyD88 single KO were as compromised as the DKOs (Fig 3F). We also evaluated the role of the TRIF-IRF3 pathway in controlling Cnbp nuclear translocation by monitoring nuclear translocation of Cnbp in macrophages from IRF3/7 DKO mice. Cnbp nuclear translocation proceeded normally in these cells (Fig 3G). Downstream of MyD88, IRAK1, 2 and 4 are engaged to control NFkB signaling. Similar to what we observed in MyD88-deficient macrophages, the LPS induced nuclear accumulation of Cnbp was reduced in cells lacking these IRAK kinases (Fig. 3H). IRAK2 and IRAK4 appeared to be the predominant drivers of this response since IRAK1 KO had residual Cnbp nuclear translocation. Consistent with a role for IRAKs as regulators of CNBP nuclear translocation, the inducible expression of IL12β was compromised in cells lacking these IRAKs (Suplemental Fig. 3A). Expression of IL-6 a known target gene of IRAK signaling but not IFNβ served as positive and negative controls in these assays respectively (Supplemental Fig 3B, C). Finally, downstream of the IRAK kinases, the kinase TAK1 is an important driver of TLR-MyD88 dependent signaling. We tested the effect of 5Z-7-Oxozeaenol, an inhibitor of TAK1 kinase. Cells treated with 5Z-7-Oxozeaenol had a dose dependent decrease in Cnbp nuclear translocation. These observations suggest that Cnbp nuclear translocation proceeds via a TLR-MyD88-IRAK-TAK1 pathway in cells exposed to LPS (Fig. 3I).

### Cnbp regulates IL12β via the NFkB/Rel family member c-Rel

Multiple DNA elements in the promoter of the IL-12β gene contribute to the inducible expression of IL-12β. These include DNA-binding sites for the NF-κB/Rel family of transcription factors, an ISRE element that binds IRF factors and activator protein 1 (AP1) elements that respond to MAPK (Fig 4A). To pinpoint which, if any of these pathways might be regulated by Cnbp we took advantage of reporter genes with multimerized DNA elements that bind each of these factors. In Hek293 cells, ectopic expression of Cnbp alone did not induce any of these reporter genes, although expression of STING induced both the NF κB and ISRE reporters (Fig. 4B). We next tested the ability of CNBP to turn on expression of a reporter gene driven by the IL12β proximal promoter. CNBP alone weakly induced the reporter but in the presence of either c-Rel or p65 elicited a robust induction of the reporter gene (Fig 4C). The ability of CNBP together with c-Rel or p65-to induce the IL12β reporter gene was dependent on a functional κB DNA binding element, since a reporter gene with a mutation in the IL12β -kB binding site was not induced (Fig. 4D). These observations indicate that CNBP has the capacity to synergize with c-Rel or p65 to turn on IL12β gene transcription. As described earlier, Cnbp-deficient macrophages had normal levels of TNFα, a well characterized NFκB target gene in response to multiple ligands including LPS, suggesting that NFκB activation was intact in cells lacking Cnbp. However to evaluate the requirement for Cnbp in controlling NFκB activation in LPS treated macrophages we monitored the phosphorylation and degradation of IkBα. IkBα was phosphorylated and degraded normally in cells lacking Cnbp (Fig 4E). Similarly, phosphorylation and nuclear translocation of the NFκB family member, p65 was intact between both genotypes (Fig 4E, F). We also evaluated the activation of the TBK1-IRF3 signaling pathway by monitoring TBK1 phosphorylation. Again TBK1 phosphorylation was equivalent in Cnbp^+/+^ and Cnbp^−/−^ macrophages (Supplemental Fig 4A).

**Figure 4.**
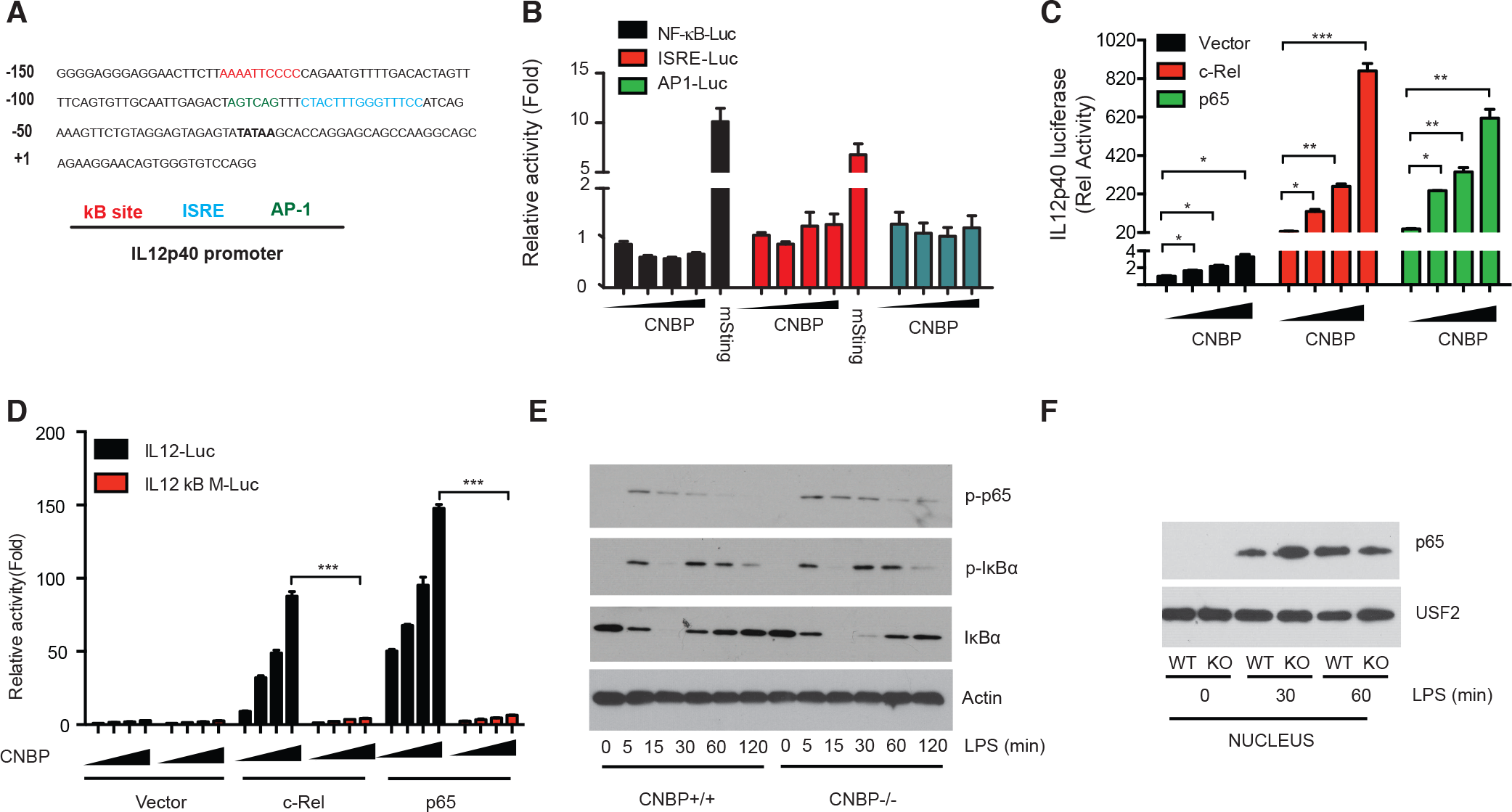
CNBP synergizes with either c-Rel or p65 to regulate IL-12p40. (A) IL12p40 promoter analysis. (B) Luciferase activity of NF-κB, ISRE or AP1 in Hek293T cells after 36-h transfection with increasing amounts of plasmids encoding Cnbp or mSting (C) Luciferase activity of IL12 p40 in Hek293T cells after 36-h co-transfection of c-Rel or p65 with increasing amounts of Cnbp (D) Luciferase activity of IL12 p40 or IL12 p40 containing κB sites mutation in Hek 293T cells after 36-h co-transfection of c-Rel or p65 with increasing amounts of Cnbp. (E) Immunoblot analysis of p-p65, p-IκBα or total IκBα in whole-cell lysates of Cnbp WT or KO BMDMs stimulated for different times with LPS (F) Nuclear extracts were analyzed for p65 by Western blotting in CNBP WT and KO BMDMs treated with LPS. Error bars represent s.e.m. of triplicate biological replicates (B-D). Data are representative of three independent experiments. *p < 0.05, **p < 0.01, ***p < 0.001. See also Figure S4.

Extensive work has revealed the importance of a c-Rel-containing NF κB complex for activation of the endogenous IL12β gene. Given the relatively specific effect of Cnbp-deficiency for IL12β gene expression and the normal activation of NFκB/p65 signaling in Cnbp-deficient macrophages we hypothesized that Cnbp might regulate c-Rel. To address this directly, we took three different approaches. Firstly, we used CRISPr/Cas9 to generate Hek293 cells lacking Cnbp and evaluated the ability of c-Rel to turn on the IL12β promoter driven reporter gene in these cells. Ectopic expression of c-Rel induced the reporter gene in Hek293 cells and this effect was impaired in Hek293 cells lacking Cbbp (Fig. 5A). We next tested if Cnbp could interact with c-Rel in macrophages. An endogenous co-immunoprecipitation (co-IP) experiment was performed and the result showed that Cnbp associated with c-Rel at steady state and that this interaction was enhanced following LPS treatment of BMDMs (Fig. 5B). Third, because c-Rel nuclear translocation is required for IL12β gene induction, we examined c-Rel nuclear translocation in Cnbp*^+/+^* and Cnbp^−/−^ macrophages. As shown in Fig. 5C, c-Rel translocated to the nucleus following LPS stimulation in Cnbp*^+/+^* macrophages but this was severely compromised in Cnbp^−/−^ macrophages (Fig 5C). In line with this finding, ChIP assay demonstrated that the LPS-induced binding of c-Rel to the IL12β promoter was greatly reduced in Cnbp-deficient BMDMs (Fig. 5D). We also evaluated the ability of Cnbp to bind the IL12β promoter. ChIP analysis revealed that Cnbp also binds the IL12β proximal reporter (Fig. 5E). Genome wide transcriptomics studies have defined the genes induced in macrophages following LPS that are dependent on c-Rel (Steve Smale, personal communication). These include IL12β as well as IL4i1 and Med21. Consistent with a specific effect of Cnbp on c-Rel activity, the induction of IL4i1 and Med21 were also compromised in cells from Cnbp-deficient mice (Fig. 5F and G). Together, these data indicate that Cnbp regulates c-Rel nuclear translocation and binding to the IL12β promoter to turn on c-Rel target genes.

**Figure 5.**
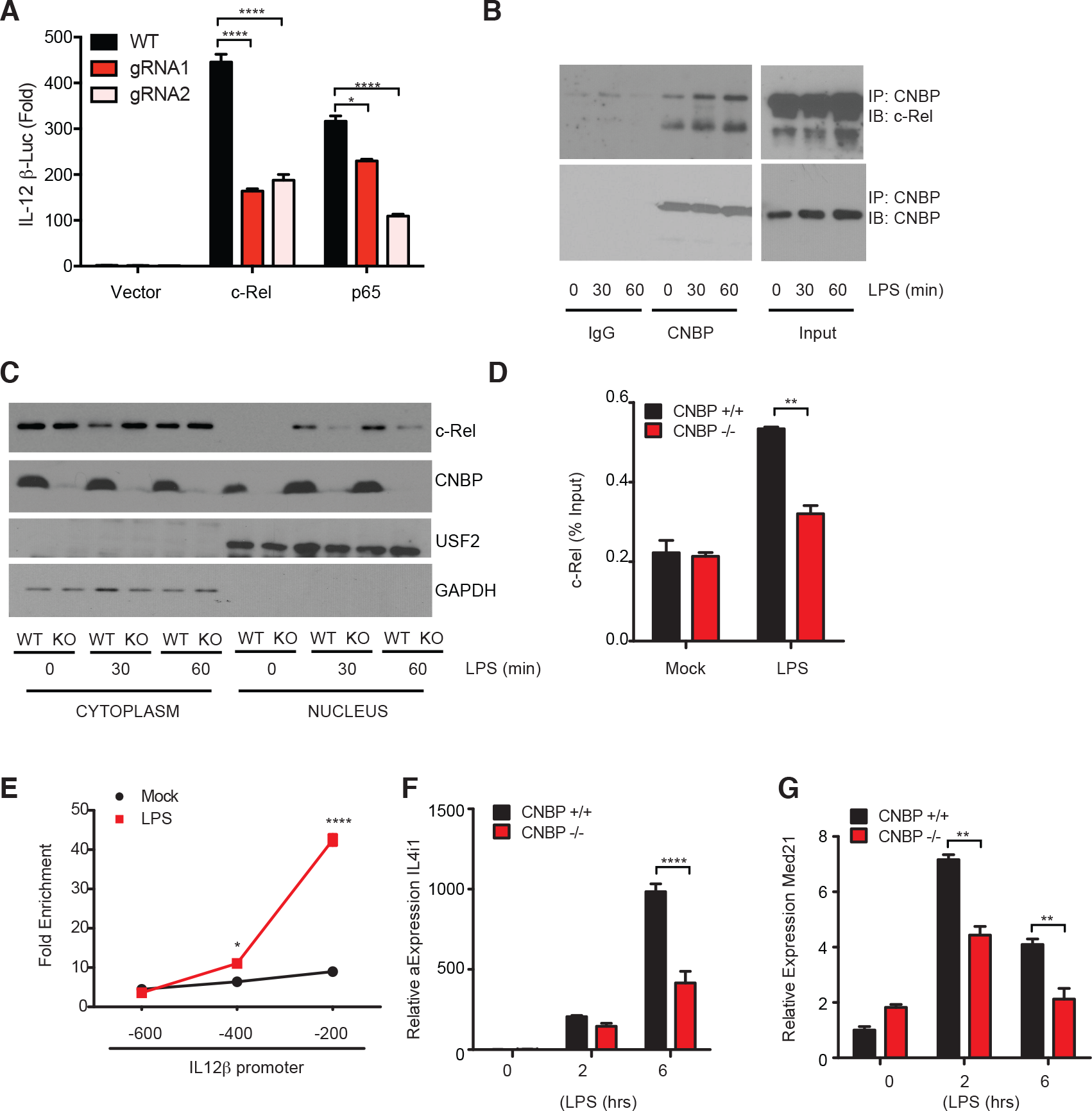
Cnbp regulates c-Rel function to control IL12p40 gene transcription. (A) Luciferase activity of IL12 p40 in Hek293T Cnbp WT or KO cells after 36-h transfection of c-Rel or p65. (B) Co-immunoprecipitation of Cnbp and c-Rel in LPS-stimulated BMDMs. (C) c-Rel nuclear translocation analysis in Cnbp WT and KO BMDMs stimulated with LPS. (D) ChIP followed by quantitative PCR (ChIP-qPCR) of c-Rel at the Il12 p40 promoter in Cnbp WT and KO BMDMs. (E) ChIP followed by quantitative PCR (ChIP-qPCR) of Cnbp at the Il12 p40 promoter in WT BMDMs (F and G) qRT-PCR analysis of Il4i1(F) and Med21(G) in Cnbp WT or KO BMDMs unstimulated or stimulated with LPS. Error bars represent s.e.m. of triplicate biological replicates (A) or s.d. of triplicate technical replicates (D-G). Data are representative of two (B and C) or three (A, D-G) independent experiments with similar results. *p < 0.05, **p < 0.01, ****p < 0.0001.

### Cnbp Protects Mice Against Infection with *T. gondii*

We next evaluated the contribution of Cnbp in controlling IL12β gene transcription and IL12β production in macrophages infected with *Mycobacterium tuberculosis, Salmonella typhimurium* and *Toxoplasma gondii*. In all cases the inducible expression of IL12β and production of IL12β protein was impaired in macrophages from *Cnbp^−/−^* mice (Fig. 6A and B). In the case of *T. gondii* infection, DCs are thought to be important producers of IL12 in vivo. We therefore purified splenic DCs and exposed them to *T. gondii* infection. Wild type and *Cnbp^−/−^* mice were previously treated with Flt3L to expand the DC pool. Isolated CD11c+ DCs were then infected in vitro with *T. gondii* and the levels of IL12β measured by ELISA. As shown in figure 6C, the levels of IL12 produced from *T. gondii* infected DCs was also reduced in the absence of Cnbp. These observations indicate that both macrophages and DCs produce IL12 in a Cnbp-dependent manner.

**Figure 6.**
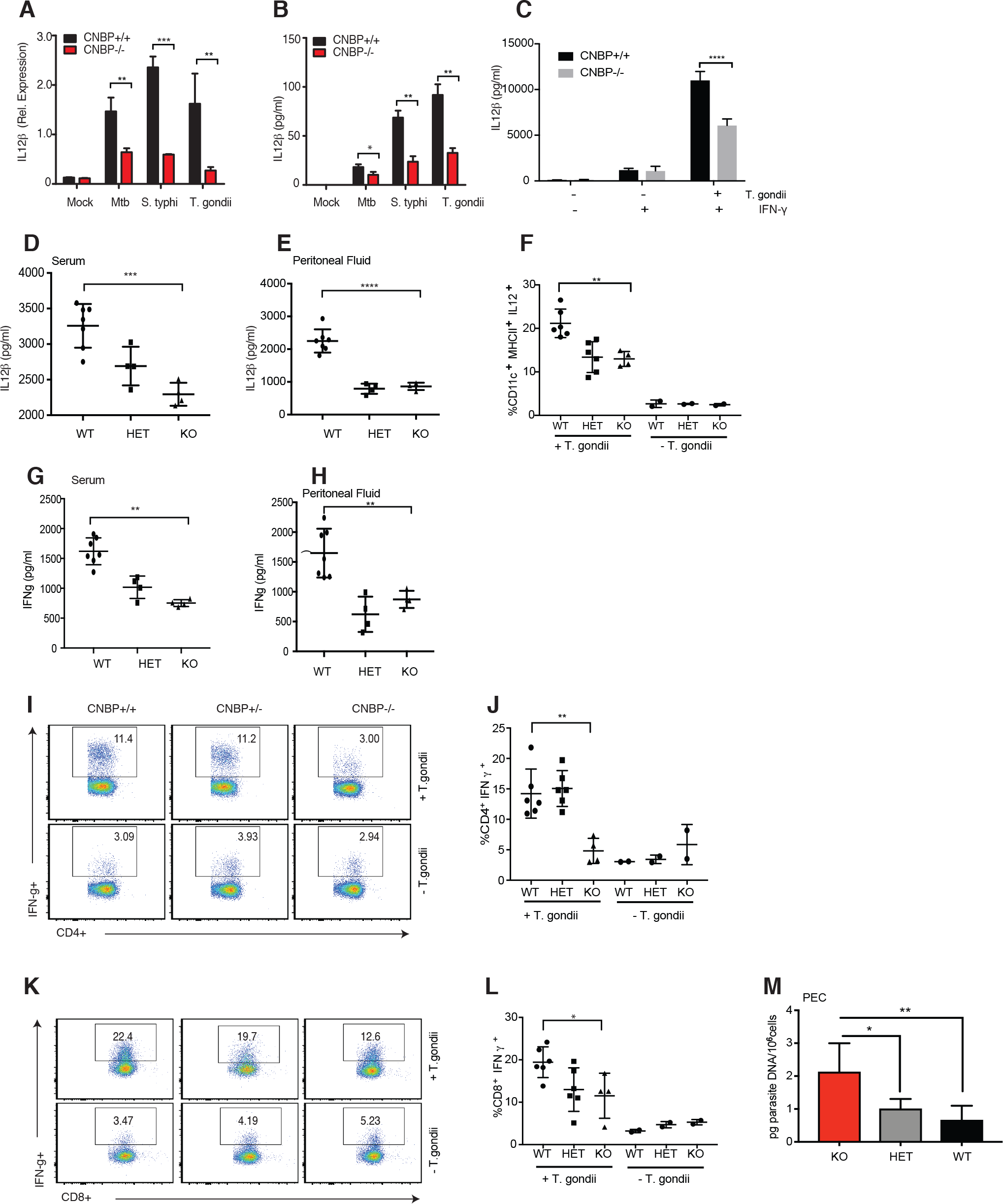
Cnbp Protects Mice Against Infection with Toxoplasma gondii. (A and B) IL12β mRNA and protein level in WT and CNBP−/− uninfected (0 h) or infected with Mtb, Salmonella or *T. gondii* were detected through qRT-PCR (A) and ELISA (B) (C) CD11c^+^ Splenocytes isolated by magnetic purification from mice injected with Flt3L were infected with *T. gondii* and the supernatant was collected for IL12β determination by ELISA (D and E) ELISA quantification of IL-12 β levels in the serum (D) and Peritoneal fluid (E) 7 d after *T. gondii* infection in WT, CNBP+/− (HET) and CNBP−/− (KO). (F) Frequency of IL-12p40 producing DCs (CD11c^+^MHC II^+^) in WT, HET and KO mice. Data are representative of one experiment (n ≥ 4) that was performed once. (G and H) ELISA quantification of IFN-γ levels in the serum (G) and Peritoneal fluid (H) 7 d after *T. gondii* infection in WT, CNBP+/− (HET) and CNBP−/− (KO). (I-L) Splenocytes were stimulated for 4 h with PMA/ Ionomycin in the presence of brefeldin A. Frequency of IFN-γ producing CD4+ (I-J) and CD8+ (K-L) T cells was measured by flow cytometry. Flow plots are gated on CD3^+^CD4^+^ (I) or CD3^+^CD8^+^ (K) T cells. (M) Parasite burden in peritoneal macrophages (PEC) at day 7 postinfection, was determined by qRT-PCR. Each symbol (D–L) represents an individual mouse; small horizontal lines indicate the mean. Data are representative of two independent experiments with similar results. *p < 0.05, **p < 0.01, ***p < 0.001, ****p < 0.0001. See also Figure S5.

The ability of IL12 to control T-lymphocyte responses leading to the production of interferon-γ is well established. Previous studies have shown that IFNγ produced by both CD4+ and CD8+ T cells plays an essential role in the protective response to *T. gondii* in vivo. To evaluate the impact of Cnbp deficiency on IL12 production, IFNγ production and protective CD4 and CD8 T cell responses *in vivo, Cnbp^+/+^, Cnbp^+/−^* and *Cnbp^−/−^* mice (generated from intercrossing heterozygous parents) were infected with the *T. gondii* (strain ME49). Spleen size and weight increased as infection progressed in all mice as expected (Supplemental Fig 5A, B, C). Consistent with our *in vitro* findings, the levels of IL12β in the serum and peritoneal fluid was greatly reduced in *Cnbp^−/−^* mice relative to Cnbp^+/+^ mice (Fig. 6D and E). Interestingly, there was also a decreased cytokine response in heterozygous mice. IL12 producing CD11c+ DCs were also reduced in *Cnbp^+/−^* and *Cnbp^−/−^* mice relative to their wild type littermate controls (Fig 6F). We also analyzed IL12 production from purified DC subsets in the spleen. All DC subsets from Cnbp-deficient mice; namely CD8^+^ DC, CD8^-^DC and moDC produced less IL12 *in vivo* (Supplemental Fig 5D).

Levels of IFNγ in the serum and peritoneal fluid of mice challenged with *T. gondii* were also reduced in *Cnbp^+/−^* and *Cnbp^−/−^* mice relative to their wild type counterparts (Fig. 6G and H). Next, we examined whether the absence of Cnbp during *T gondii* infection altered the CD4 and CD8 T cell response. T cells from *Cnbp^+/+^, Cnbp^+/−^* and *Cnbp^−/−^*infected mice were stimulated ex vivo with PMA and ionomycin and IFNγ production from CD4+ and CD8+ T cells measured (Fig. 6I). The percentage of IFNγ positive CD4 T cells was significantly reduced in *Cnbp^−/−^* mice (Fig. 6J). Similarly, the percentage of IFNγ positive CD8 T cells was also lower in these animals (Fig. 6K and L). Lastly, we measured parasite loads in these animals. Cnbp-deficient mice had an increased parasite burden (Fig 6M). Altogether, these results indicate that Cnbp-deficient mice fail to mount protective IL-12 and IFNγ responses *in vivo*, resulting in a reduced Th1 cell immune response and an inability to control parasite replication.

## Discussion

The NFkB/Rel family of transcription factors are major regulators of the innate and adaptive immune response (Silverman and Maniatis, 2001; Zhang and Ghosh, 2001). Active p65/p50 dimers are found in the nucleus of macrophages following ligation of multiple classes of PRRs including Toll-like receptors, C-type lectin receptors, RIG-I like receptors and DNA sensors. NF κB DNA binding elements in the promoters of immune genes are bound by these complexes leading to the inducible expression of hundreds of cytokines and anti-microbial molecules in temporally distinct manners(Caamano and Hunter, 2002; Dev et al., 2011). The c-Rel/NFκB family member appears to be unique in its ability to activate transcription of the IL12β gene. The NF κB binding sites in the promoter of the IL12β gene display comparable binding affinity for both p65/p50 and c-Rel/p50 heterodimeric complexes (Sanjabi et al., 2000). Despite this, however, only c-Rel has been shown to be essential for IL12β production in macrophages stimulated through multiple TLRs and in cells infected with intracellular pathogens. These findings therefore implicate c-Rel as an essential and specific regulator of IL12β gene expression, distinguishing c-Rel from the more broadly acting RelA (Mason et al., 2002; Sanjabi et al., 2000). While the molecular mechanisms involved in c-Rel binding to the IL12β promoter and c-Rel dependent transcription of IL12β have been studied extensively, it is still unclear what governs the highly specific effects of c-Rel for IL12β regulation.

In this study, we define Cnbp as a new regulator of this c-Rel-dependent IL12β response. We identified Cnbp in dsDNA pulldown experiments that were designed to identify cytosolic nucleic acid sensors. RNA sensors such as RIG-I and DNA sensors such as cGAS and AIM2 are activated when nucleic acids gain access to the cytosol during infection leading to protective anti-microbial defense mechanisms (Thompson et al., 2011). In an effort to broaden our understanding of the DNA sensing pathway in particular, we used Mass Spectrometry to identify cytosolic DNA binding proteins and leveraged loss of function approaches to link candidate proteins to innate immunity in macrophages. We generated mice lacking Cnbp and investigated type I interferon induction in response to dsDNA viruses, polydAdT and ISD, (synthetic dsDNAs) and cyclic dinucleotides. The hallmark response of this pathway namely the type IFN response was intact in cells lacking Cnbp as was transcription of the NF κB regulated cytokine TNFα. However, we found that cells lacking Cnbp were compromised in their ability to turn on transcription of IL12β. The importance of Cnbp for IL12β gene transcription was also observed with LPS, one of the most potent inducers of IL12β expression, as well as polyI:C and Sendai virus which signal via MDA-5 and RIG-I, respectively.

In response to multiple ligands, Cnbp translocated to the nucleus and in the case of LPS, this proceeded in a manner dependent on MyD88, IRAK4, IRAK1 as well as TAK1 kinase. It is likely that for different PRRs, for example Sendai virus signaling via RIG-I, the pathway components involved would be different. Multiple genetic elements have been defined in the IL12β promoter including those that bind NF κB, IRFs (ISRE) or AP1. Overexpression of Cnbp did not induce reporters driven by κB, ISRE or AP1, however Cnbp could induce expression of a reporter gene consisting of the IL12 promoter, an effect that was greatly enhanced by co-expression of either c-Rel of RelA(p65). The κB binding element was essential for this effect since mutation of this site abrogated this response. A recent study identified Cnbp as a regulator of NF κB activation controlling sustained expression of IL-6 and a broad panel of additional cytokines in macrophages (Lee et al., 2017). In our study, however using a clean genetic approach, we found that the activation of NF κB assessed by measuring IκBa phosphorylation and degradation as well as RelA(p65) nuclear translocation) proceeded normally in cells lacking Cnbp. In contrast, we found that c-Rel was the major target of Cnbp. Firstly, c-Rel dependent activation of the IL12 promoter-driven reporter gene was impaired in HEK293T cells generated to lack Cnbp. Secondly, we found that Cnbp interacted with c-Rel and finally that c-Rel nuclear translocation and DNA binding to the IL12β promoter using Chromatin Immunoprecipiation after LPS treatment were all dependent on Cnbp. ChIP analysis in primary BMDMs also revealed binding of Cnbp to the IL12β promoter. We observed very strong binding of Cnbp to the −200 site within the IL12β promoter, where the κB binding sites reside after LPS treatment. Whether this is direct binding of Cnbp to the IL12 promoter or perhaps only because of its association with c-Rel remains unclear at present. Our data supports a model whereby Cnbp controls c-Rel nuclear translocation. It’s possible that a stable c-Rel:CNBP complex is formed in the cytoplasm and upon LPS stimulation, this complex translocates to the nucleus. Another possibility is that the c-Rel:CNBP complexes are essential for retention of c-Rel in the nucleus after inducible translocation and DNA binding. A third possibility is that CNBP influences the nuclear translocation of c-Rel by indirect mechanisms. Consistent with this latter possibility, although Cnbp and c-Rel interact, only a modest fraction of c-Rel is stably associated with CNBP. Extensive work from Smale and colleagues have defined the importance of c-Rel in controlling IL12β gene transcription (Sanjabi et al., 2000). Surprisingly, c-Rel regulates only a very restricted set of target genes. RNA-seq experiments comparing WT and c-Rel deficient macrophages stimulated with lipid A show that IL12β and a limited set of genes including IL4i1 and Med21 represent bona fide c-Rel targets. Consistently, in our experiments both IL4i1 and Med21 expression were also decreased in Cnbp-deficient macrophages stimulated with LPS. Together all of these findings implicate Cnbp as a highly specific regulator of c-Rel and define a novel TLR-Cnbp-c-Rel signaling axis controlling IL12β expression. Our studies also reveal that Cnbp nuclear translocation and regulation of IL12β is a new signaling event downstream of multiple PRRs.

Classically, IL-12 is induced by TLRs in response to microbial products (Andrade et al., 2013; Bafica et al., 2005; Schamber-Reis et al., 2013; Seki et al., 2002). Cnbp was essential for IL12β production in macrophages stimulated with any ligand tested or microbial infection. This included viruses, *M. tuberculosis, S. typhimurium* and *T. gondii* suggesting that IL12β induction regardless of the stimuli required Cnbp. Moreover, we provide important *in vivo* evidence for the importance of Cnbp in the immune system. *T. gondii* is an intracellular protozoan parasite whose elimination requires production of IFNγ that activates various cell intrinsic antiparasitic defense pathways within infected cells (Yap and Sher, 1999). The importance of IL12 in induction T-lymphocyte production of IFNγ and resistance to *T. gondii* and other intracellular pathogens *in vivo* is well established(Jordan et al., 2010; Mason et al., 2004). Consistent with the important role of Cnbp in controlling innate production of IL-12 in vitro, Cnbp-deficient mice have reduced IL12β levels in vivo. This leads to a reduction in IFNγ produced by both CD4+ and CD8+ T cells and an inability to control *T. gondii* growth *in vivo*. CD8a+ DCs and monocyte derived DCs are the important producers of IL-12 in vivo during *T. gondii* infection (Dupont et al., 2015; Goldszmid et al., 2012; Mashayekhi et al., 2011). In our studies, we found that DCs from Cnbp-deficient mice had reduced IL12 levels in vivo. Both CD8α+ DCs and CD8α− DCs produced IL-12 in a Cnbp-dependent manner. Consistently, IFNγ production by CD4 T and CD8 T cells was impaired and Cnbp knockout mice infected with *T. gondii* infection. These observations indicate that both macrophages and DCs produce IL12 in a Cnbp-dependent manner, with the latter being more relevant in vivo for development of Th1 immunity and control of *T. gondii in vivo*.

Prior to this work linking Cnbp to the IL12β and Th1 immune response, Cnbp was shown to regulate forebrain development by controlling cell proliferation and apoptosis (Calcaterra et al., 2010). It is worth noting here that we Cnbp-deficient mice were viable and in terms of their immune system were comparable to wild type mice under steady state conditions. In addition to these effects, Cnbp has been implicated in the transcriptional and posttranscriptional control of gene expression. Early studies identified Cnbp as a regulator of the transcription of c-myc, wnt and skeletal muscle chloride channel 1 (clc1) (Chen et al., 2013; Margarit et al., 2014). Recently, a PAR-CLIP study indicated that CNBP preferentially bound G-rich elements in mRNA coding sequences increasing their translational efficiency (Benhalevy et al., 2017). There is little evidence linking Cnbp to the immune system. Cnbp has been implicated in myotonic dystrophy type 2 (DM2) and sporadic inclusion body myositis (sIBM) both of which are associated with increased autoantibodies suggesting some links to the immune response (Niedowicz et al., 2010; Tieleman et al., 2009). In macrophages the ability of Cnbp to regulate NFkB and sustained expression of IL-6 was the first evidence of an immune role for this factor (Lee et al., 2017). Consistent with the effects we describe herein, this study also demonstrated using a xebrafish model that cnbp-deficient zebrafish were more susceptible to infection with *Shigella flexneri* infection suggesting that conservation of the immune regulatory effects we describe.

In conclusion, our findings provide critical new insights into the regulation of the innate immune response in macrophages and dendritic cells, describing a previously unknown Cnbp-cRel signaling axis important in controlling IL12b expression downstream of multiple PRRs. This pathway also has important effects *in vivo* controlling Th1 immunity and resistance to the parasitic pathogen *Toxoplasma gondii*. Given the importance of IL12 in control of Th1 immunity and the potential for dysregulation of the IL12-Th1 response in inflammatory diseases such as Inflammatory Bowel Disease, the identification of Cnbp as a specific regulator of c-Rel dependent IL12 induction has enormous implications for both host-defense and inflammatory disease. IL-12 is a prominent target for clinical intervention in the treatment of inflammatory diseases and cancer. Our work unveils a new target Cnbp which may provide a unique handle to specifically curtail this IL12 response.

## Acknowledgments

We are grateful to Dr. Steve Smale for providing the reporter plasmid Rel Mut p40-Luc.We would like to thank Roland Elling for providing mouse tissue RNA samples, Natalia Martinez Araujo for help with T. gondii infections and all members of the Fitzgerald laboratory for their helpful comments. This study was supported by NIH grants R37-AI067497 (to K.A.F.), by R01-AI116577 and R01-NS098747 (to R.T.G.), by R01-AI106934 (to D.M.K.).

## Author Contributions

K.A.F. supervised the work. Y.C. and K.A.F. designed the research, analyzed results, and wrote the manuscript. Y.C. performed the majority of the experiments, with contributions from S.S., P.A.A., Z.J., A. J. O., S.H., J.B., J.R.H., C.M.S. and R.T.G.. D.M.K. provided critical reagents and suggestions. All authors read and provided suggestions during the preparation of the manuscript.

## STAR Methods

### Generation of CNBP Knockout Mice

CNBP-deficient mice were generated using ES cells obtained from the Knockout Mouse Repository (KOMP, UC Davis). ES cells (Cnbp^tm1a(KOMP)Wtsi^) were generated by replacing the CNBP genomic locus with a neomycin cassette under control of a Pgk1 promoter. ES cells were injected into blastocytes to generate chimeric mice at the UMMS Transgenic Core. CNBP heterozygous mice were obtained by gamete line transmission from mating the chimeric mice with WT C57BL/6 mice. CNBP heterozygous mice were intercrossed to generate wild type and KO alleles for experimentation. Littermate controls were used throughout. Myd88-/-; Myd88-/-Trif-/- mice were from S. Akira (Tokyo, Japan). Irf3-/–Irf7-/- mice were obtained from T. Taniguchi (Japan) and used as described; All mouse strains were bred and maintained under specific pathogen-free conditions in the animal facilities at the UMass Medical School in accordance with the Institutional Animal Care and Use Committee.

### Cell Culture and Stimulation

BMDMs were generated by differentiating bone-marrow cells in DMEM supplemented with 10% fetal calf serum (FCS), 1% Penicillin/Streptomycin (P/S) cocktail and 20% L929 supernatant for 7 days. BMDCs were generated by differentiating bone-marrow cells in RPMI 1640 medium supplemented with 10% FCS, 1% P/S cocktail and recombinant GM-CSF (20 ng/ml) for 10 days. For all experiments, cells were plated one day prior to stimulation. Cells were stimulated at following concentrations (unless mentioned otherwise): LPS (100 ng/ml), poly(I:C) (25 µg/ml) and BAY 11-7082 (10 µM). Transfection of BMDMs with poly(dA:dT) and ISD was performed using lipofectamine 2000 (Invitrogen). BMDMs were infected with Herpesviruses Herpes simplex virus (HSV-1-ICP0-deficient mutant) were from D. Knipe (HMS, Boston, MA) (10 MOI), mCMV (10 MOI), IAV (5 MOI), Sendai virus (200 IU/ml), *Mycobacterium tuberculosis* (10 MOI), *Salmonella* (10 MOI), *Toxoplasma gondii* (5 MOI) for indicated hours for mRNA and protein analysis.

### Parasites

The Toxoplasma gondii strain ME49 was maintained in C57BL/6 mice by serial inoculation of brain homogenate-containing cysts. ME49 tachyzoites were maintained in human foreskin fibroblast cells (Hs27). Parasites suspentions were prepared by collection of infected cells and disruption by repeated passage through a 27-gauge needle followed by centrifugation at 900g for 10 min. For STAG preparation, the parasites were submitted to 3 cycles of freezing at -70°C and thawing at 37°C followed by four 20-s rounds of sonication. The resulting homogenate was then centrifuged at 600g for 10 min, and the supernatant was collected and its protein concentration determined by the Bradford method (Invitrogen).

### In vivo experimental infections

Mice were inoculated intraperitoneally (i.p.) with 25 cysts obtained from brain homogenates of 6–8 weeks infected mice. Mice were monitored for survival or sacrificed 7 days post infection in order to collect peritoneal exudate cells and fluid, blood, liver and spleens. Samples from each mouse were individually processed and analyzed. Quantification of T. gondii DNA was performed by amplification of B1 gene using the primers (Fw: 5’- CTGGCAAATACAGGTGAAATG-3’; Rv: 5’- GTGTACTGCGAAAATGAATCC-3’) and analyzed based on a standard curve of purified parasite DNA. PCR reactions were setup in a final 20 ml volume using 5 ng of total tissue DNA, 200 ng of each primer and 1x of the iQ SYBR Green Supermix (BioRad)

### In vitro infection

Macrophages cultures were induced or not overnight with IFN-γ (40 ng/mL) before infection. Cells were infected with ME49 tachyzoites (MOI=5) and after 2h, unbound parasites were washed off and the cultures were incubated for 24h to obtain supernatant and 4h for RNA extraction.

### Affinity purification with biotinylated Oligo

Cytosolic cell extracts were generated from 10x106 murine bone marrow derived macrophages or 20x106 THP1 cells using low salt lysis buffers*. The lysates were precleared with Streptavidin (SA) ultralink agarose protein G resin for 2h at 4°C and the resin washed out. 3µg of Biotinylated oligos were then added to precleared lysates for binding overnight (~18h) at 4°C. SA ultralink agarose G-beads were added to bind biotinylated complexes for 2h at 4°C. The bound complexes were pelleted by spinning at 6500 RPM for 5 min at 4°C. Several rounds of increasingly stringent washes were carried out to reduce non-specific binding partners. The majority of the sample was run under non-denaturing conditions on a stacking gel composition PAGE. A shotgun approach was used to run the sample i.e. the sample was run so that all of it had migrated into the gel. The gel was subsequently stained with MS compatible Biosafe Coomassie blue (Biorad) to identify the band of proteins in the gel and the band was excised.

A small aliquot was subjected to normal SDS page gel and western blotting for QC.

### LC-MS/MS

Isolated proteins in the complex were identified and analyzed by LC/MS at the Mass spectrometry Core Facility at UMass. Briefly, DNA bound protein (≥2pmol of protein) was centrifuged and subsequent pellet vacuum dried. The pellet was then dissolved in 20ul of 0.1% Rapigest detergent in 0.1M tetramethylammonium bicarbonate. Disulfides were reduced with DTT for 30min at 60°C. Sulfhydrol were alkylated with 20nmoles of iodoacetamide for 3h in the dark at 22°C. Trypsin digestion and capillary LC-MS/MS were performed on the ThermoFinnigan LTQ quadrupole ion trap system. Proteins were identified from the product ion scan spectra using SEQUEST search engine with species subset of the IPI protein database. Peptide and protein data was assembled in the Scaffold software and analyzed against proteins bound to control DNA ODN. Proteins were subsequently identified by their GO annotations and sorted through the domain database at Pfaam (Wellcome Trust, Sanger Institute) for initiat bioinformatics predictions.

### RNA extraction and RT-qPCR

Total RNA from BMDMs, BMDC and peritoneal exudate cells was extracted with RNeasy RNA extraction kit (Qiagen) according to the manufacturer’s instructions. Genomic DNA in RNA purifications were eliminated using on-column DNase I digestion (Qiagen). Equal amounts of RNA (1000 ng) was reverse-transcribed using iScript cDNA synthesis kit (Bio-Rad). Diluted cDNAs (1: 100 final) were subjected to qPCR analysis using iQ SYBR Green Supermix reagent (Bio-Rad). Gene expression levels were normalized to Gapdh as housekeeping genes. Relative mRNA expressions were calculated by the change-in-cycling-threshold method as 2-ddC(t). The specificity of amplification was assessed for each sample by melting curve analysis. Primer sequences are provided in supplemental.

### Cytokine analysis

Cell culture supernatant were assayed for cytokine levels using commercially available sandwich ELISA kits: IL12p40, IL10, IFN-γ (R&D Systems), IL6 (BD Biosciences) and TNFα (eBioscience). the murine IFN-β kit is as described (Roberts et al., 2007). All experiments for cytokine analysis by ELISA were performed in biological triplicates.

### NanoString analysis

Cell stimulation and RNA isolation was performed as described above. The nCounter analysis system was used for multiplex mRNA measurements using the custom gene expression code-set against 200 immune genes. Total RNA (100 ng) was hybridized overnight with the gene expression code-set and analyzed on an nCounter Digital Analyzer (Nanostring Technologies). RNA hybridization, data acquisition and analysis was performed as per manufacturer’s specifications. RNA counts were processed to account for hybridization efficiency, and mRNA expressions across experimental groups were normalized to the geometric mean of six housekeeping genes

### Chromatin Immunoprecipitation (ChIP)

ChIP experiments were performed essentially as described(Atianand et al., 2016). Briefly, 1 *107 primary BMDMs were used to perform immunoprecipitation with mouse monoclonal anti-Cnbp. qPCR was performed on immunoprecipitated and input fractions from the immunoprecipitation.

### Purification of CD11c+ DCs

Mouse CD11c+ DCs were purified from spleens of mice injected with BL6-FLT3L producing cells. Briefly, mice were injected subcutaneously with 2 × 106 cells in 300µl PBS. Spleens were harvested within 2 weeks after injection and CD11c+ DCs were isolated using the CD11c MicroBeads UltraPure (Miltenyi Biotec). B16-FLT3L producing cells were from G. Dranoff (HMS, Boston, MA)

### Flow cytometry

Mononuclear cells were incubated with 50 ng/ml phorbol myristate acetate (PMA) (Sigma) and 500 ng/ml ionomycin (Sigma) in the presence of GolgiStop (BD) in complete T cell media at 37°C for 4 h. Cells were washed with FACS buffer, stained with Live/Dead fixable aqua dead cell marker (Invitrogen) and incubated with Fc block prior to staining. The following Abs were used for staining: PerCP-Cy5-CD8; APC-Cy7-CD8; APC-F4/80; Alexa700-CD11c; FITC-MHCIIb; PE-Cy7-CD11b; eFlour 450-Ly6C; APC-Cy7-CD4; FITC-CD3; Intracellular staining for APC-IL4; PE-Cy7-IL17a; eFluor450-IFNg; PE-IL12. Intracellular cytokine staining was performed by fixing cells in 2% paraformaldehyde, followed by permeablization and staining (BD Biosciences). Flow cytometric analyses were performed on an LSRFortessa (BD Biosciences) and analyzed using FlowJo Software (Tree Star).

### Immunofluorescence staining

Primary BMDMs were fixed in 4% paraformaldehyde, permeabilized with 0.2% Triton X-100/PBS before incubation with primary antibodies (anti-Cnbp) for 2 hour at room temperature (RT). Cells were washed in PBS, followed by incubation with TRITC-conjugated anti-mouse immunoglobulin (Ig) secondary antibodies. Nuclei were stained with DAPI.

### luciferase reporter assay

Hek293T cells were seeded on 96-well plates (4 × 104 cells per well) and then transfected with 50 ng IL12p40, IL6 or TNF–luciferase reporter vector and 5 ng renilla-luciferase reporter vector with increasing amounts of expression vector for Cnbp or plus 10 ng c-Rel or p65 expression vector. Empty control vector was added so that a total of 200 ng vector DNA was transfected into each well. Cells were collected at 36 h after transfection and luciferase activity was measured with a Dual-Luciferase Reporter Assay System according to the manufacturer’s instructions (Promega). Firefly luciferase activities were normalized to Renilla luciferase activities. All reporter assays were repeated at least three times. Data shown were average values and SEM from one representative experiment.

### Co-immunoprecipitation and Western Blot Analysis

Cells were lysed 36–48 h after transfection of expression plasmids using 50 mM Tris–HCl, pH 8.0, 150 mM NaCl, 1% Triton X-100 containing cocktail. For immunoprecipitation, lysates were incubated with the appropriate antibodies for 2 h on ice, followed by precipitation with protein G Sepharose. Samples were separated by SDS–PAGE and transferred to PVDF membranes. After blocking in PBS containing 0.1% Tween-20 and 5% skim-milk, the blots were probed with indicated antibodies. Westernblot visualization was done with enhanced chemiluminescence.

### Statistics

GraphPad Prism 5 software (GraphPad Software, San Diego, CA) was used for data analysis using a two-tail unpaired t test. A p value, 0.05 was considered statistically significant (*p <0.05, **p, <0.01, ***p < 0.001, ****p < 0.0001).

**Figure S1:**
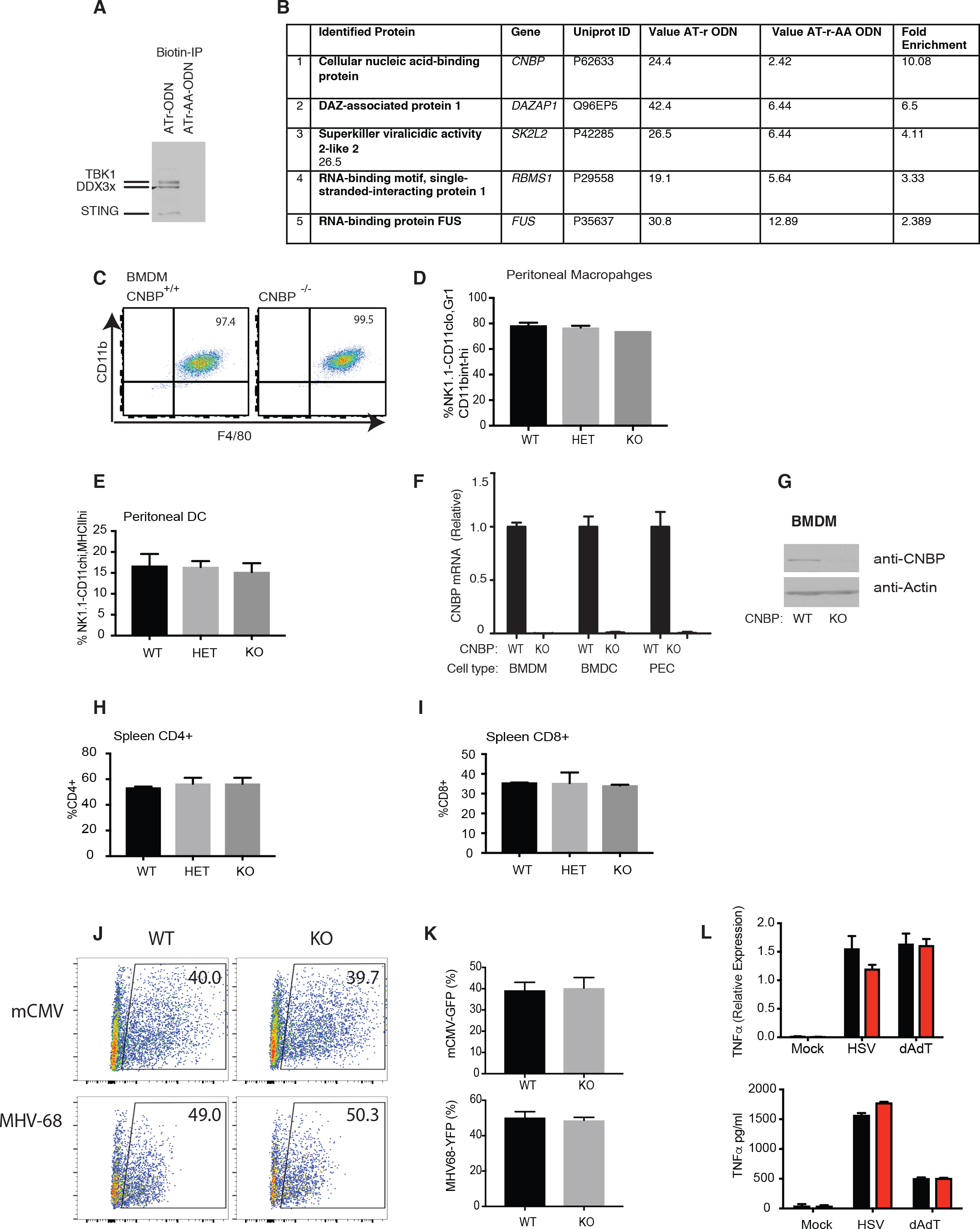
Characterization of CNBP deficient mice. **Related to Figure 1** (A) Immunoblotting was performed to identify known components of DNA sensing pathway in ATr-ODN and a non-stimulatory ODN (ATr-AA ODN) pulldowns from macrophage cytosolic extracts. (B) LC-MS analysis of proteins identified. The top 5 most enriched proteins identified in ATr-ODN and non-stimulatory ATr-AA-ODN complexes. (C) CD11b/F480 levels measured by flow cytometry on macrophage cultures differentiated in vitro from bone marrow progenitors. (D and E) Percentages of peritoneal macrophages (D), peritoneal DC (E) were analyzed by their specific lineage markers from control and CNBP-deficient mice. (F and G) CNBP expression level in WT and KO cells were detected through RT-PCR (F) and Western blotting (G). (H and I) Percentages of CD4 and CD8 T cells analyzed by their specific lineage markers from control and CNBP-deficient mice. (J and K) The replication of reporter virus mCMV-GFP or MHV-68-YFP in CNBP WT or deficient BMDMs. (L) TNFα mRNA and protein level in CNBP+/+ and CNBP−/− BMDMs treated with HSV or dAdT were detected through qRT-PCR and ELISA.

**Figure S2:**
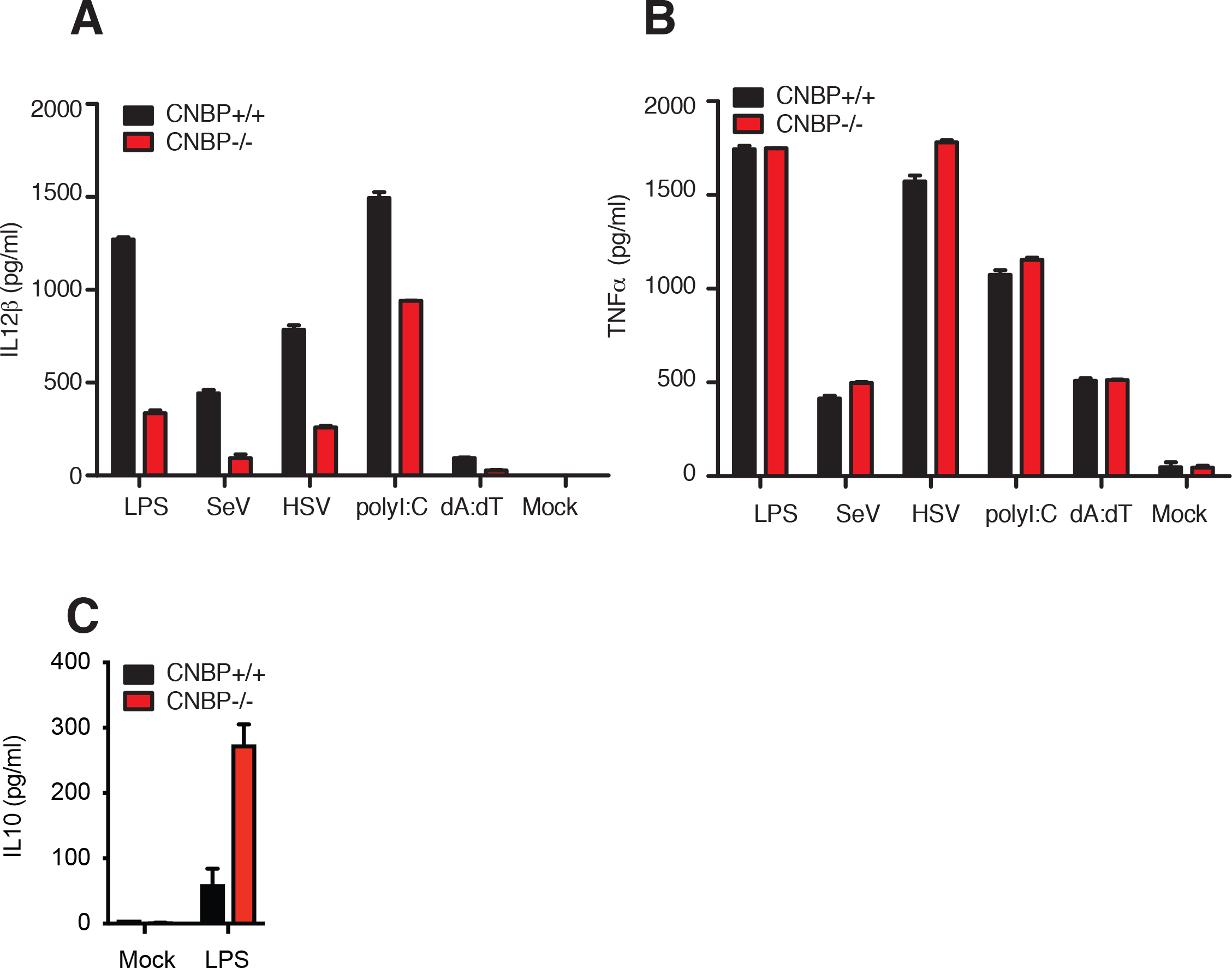
Cytokines analysis in CNBP KO cells after various ligands stimulation. **Related to Figure 2** (A and B) ELISA analysis of IL12β (A) and TNFα (B) in Cnbp WT or KO BMDMs left unstimulated or stimulated with LPS, SeV, HSV, PolyI:C or transfected dA:dT. (C) ELISA analysis of IL10 in Cnbp WT or KO BMDMs left unstimulated or stimulated with LPS.

**Figure S3:**
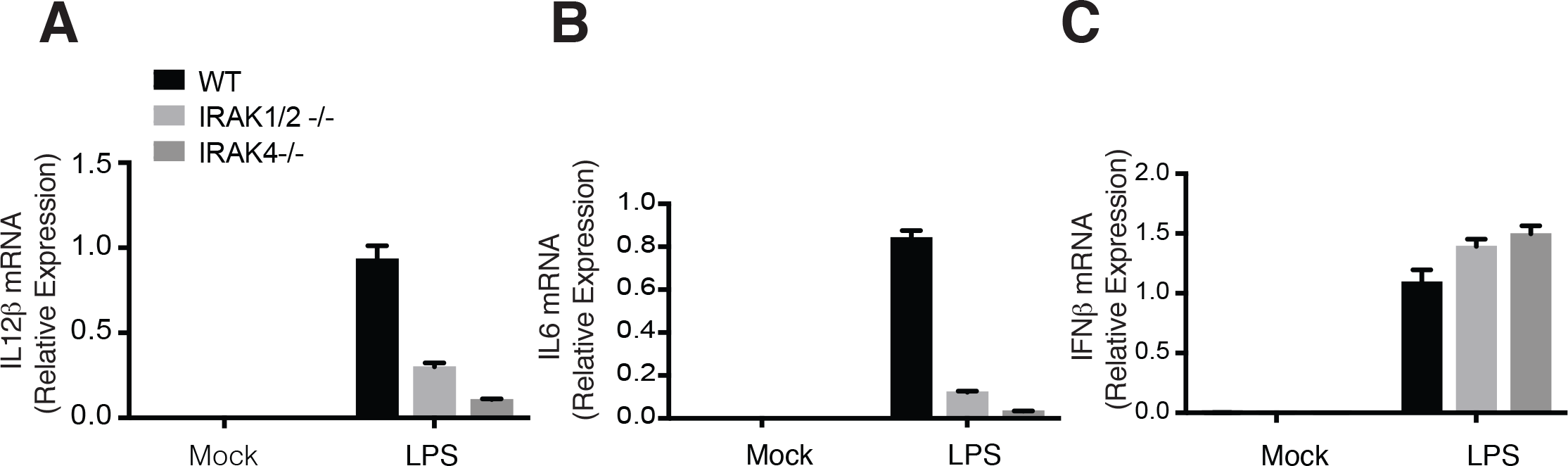
IL12 and IL6 mRNA decrease in IRAK1/2−/− or IRAK4−/− BMDMs. **Related to Figure 3** (A-C) qRT-PCR analysis of IL12β (A), IL6 (B) and IFNβ (C) in IRAK1/2-/- or IRAK4-/- BMDMs left unstimulated or stimulated with LPS.

**Figure S4:**
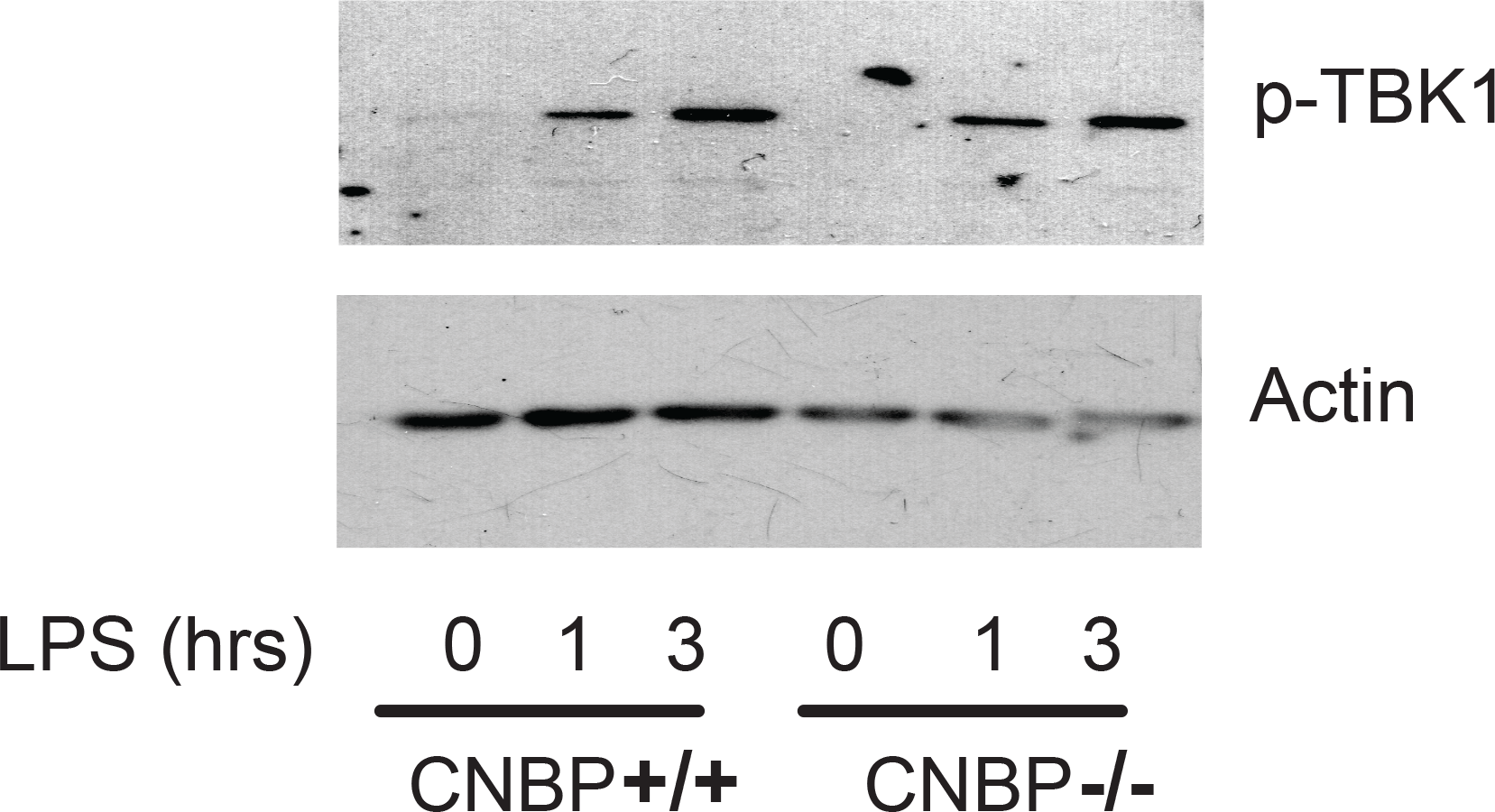
TBK1 phosphorylation was equivalent in Cnbp+/+ and Cnbp−/− macrophages. **Related to Figure 4** Immunoblot analysis of phosphorylated (p-) TBK1 in whole-cell lysates of Cnbp WT or KO BMDMs stimulated for various times with LPS.

**Figure S5:**
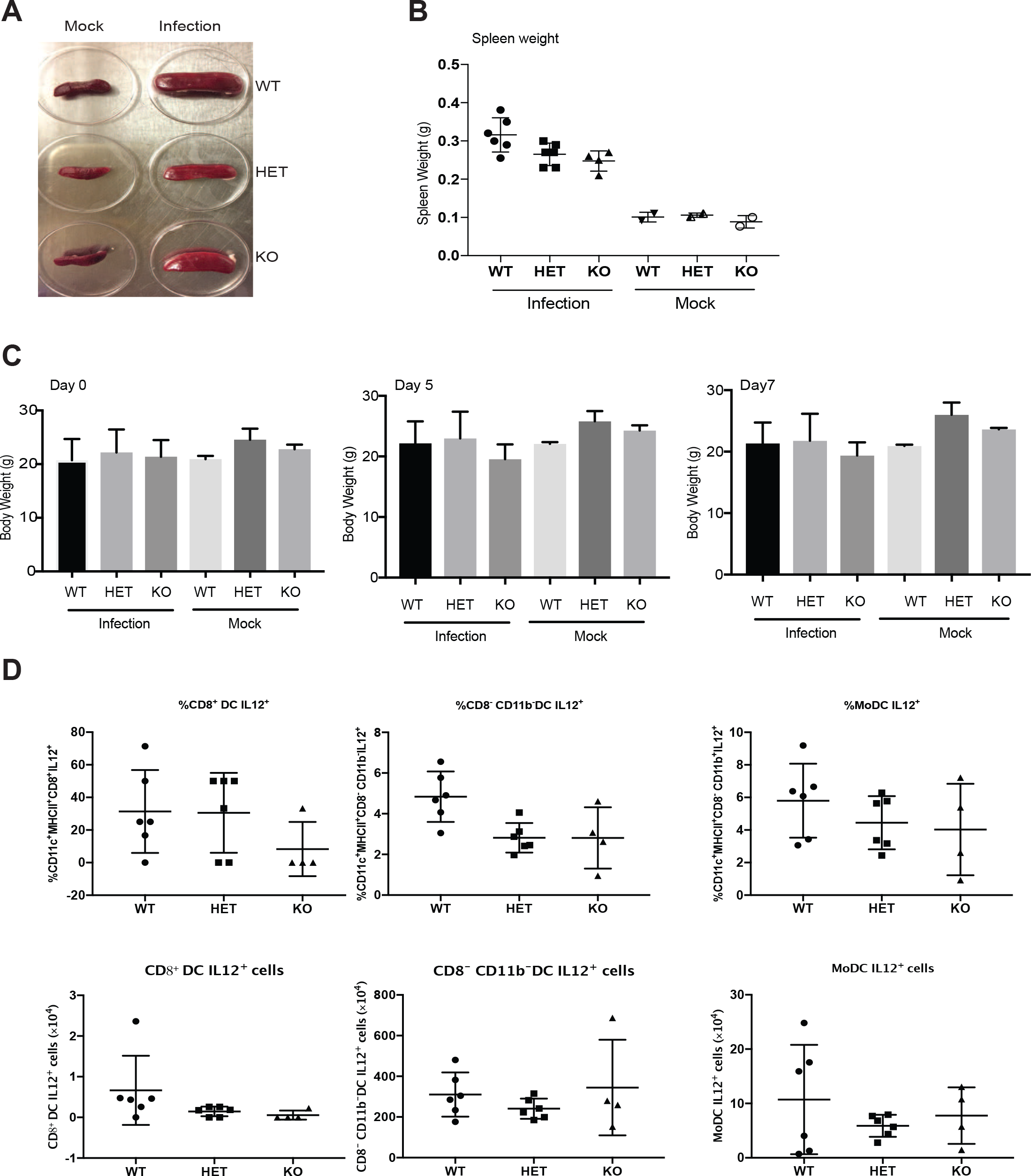
The analysis of CNBP deficient mice after *T. gondii* infection. **Related to Figure 6** (A and B) Gross appearance (A) and weight (B) of the spleen in the indicated mice (C) Body weight of the mice uninfected or infected with *T. gondii*. (D) IL12 production in different DC subsets from Cnbp-deficient mice

